# Integrating genetics, morphology, and fungal host specificity in conservation studies of a vulnerable, selfing, mycoheterotrophic orchid

**DOI:** 10.1101/2020.08.17.254078

**Authors:** Nicole M. Fama, Brandon T. Sinn, Craig F. Barrett

**Affiliations:** Department of Biology, West Virginia University, 53 Campus Drive, Morgantown, West Virginia, USA 26506; National Institutes of Health, Building 10 Room 5-3816, 10 Center Drive, Bethesda, Maryland, USA 20854; Department of Biology and Earth Science, Otterbein University, 214 Science Center, 1 South Grove Street, Westerville, Ohio, USA 43081

**Keywords:** Autogamous, cleistogamous, ISSR, rare, symbiosis

## Abstract

Mycoheterotrophic plants derive most or all carbon and nutrients from fungal partners and represent poorly understood components of forest biodiversity. Many are rare or endangered yet can be ecological indicators of forest ecosystem function due to their often highly specific fungal host requirements. One such species is the IUCN red-listed (‘vulnerable’), fully mycoheterotrophic orchid, *Corallorhiza bentleyi*. This recently described species is among the rarest plants in Appalachia, known from five counties in Virginia and West Virginia, USA. The species has a restricted range, small population size, and is self-pollinating. Here we take an integrative approach to conservation genetic assessment in *C. bentleyi* using floral morphometrics, simple-sequence repeats, and fungal host DNA to characterize variation within and among sampling localities. Morphology reveals some differentiation among individuals from six sampling localities. Surprisingly, most genetic variation is found within localities, contra to the expectation for a selfing species. Fungal host DNA reveals extreme specificity upon a few genotypes of a single ectomycorrhizal host species, *Tomentella fuscocinerea,* across all localities. We discuss the conservation implications of morphological, genetic, and symbiotic diversity in this vulnerable species, and recommend additional assessment of conservation status based on: an obligate reproductive mode of selfing, preventing benefits of outcrossing among genetically non-identical individuals; extreme host specificity, severely restricting niche space; and highly fragmented habitat under threat from anthropogenic disturbance. This study underscores the importance of integrative conservation assessment, analyzing multiple data sources, and reveals patterns not readily apparent from census-based assessments alone.

## INTRODUCTION

Mycoheterotrophs are leafless plants that parasitize mycorrhizal fungi for nutrition and remain some of the most poorly understood species in terms of population dynamics, genetic diversity, and factors controlling species distribution and abundance (Leake et al. 1994; Bidartondo 2005; Merckx et al. 2013; Taylor et al., 2002; Taylor et al., 2013; Waterman et al., 2013; Gomes and Merckx, 2019). They represent a particularly difficult element of forest ecosystem dynamics, due to their dependence upon often ephemeral or rare fungal taxa that are even more poorly understood. They have been described as ‘ecological indicators’ of forest habitat health and mycorrhizal diversity (Barrett et al., 2010; Merckx et al., 2013), thus underscoring the need to better understand their unique ecological attributes. Orchids make up the majority of all mycoheterotrophic plant species, with >200 leafless, fully mycoheterotrophic species globally (Freudenstein and Barrett, 2010; Merckx et al., 2013).

Habitat fragmentation via deforestation for agriculture, extraction industries, and development over the last several hundred years has had profound, complex, and global effects on genetic diversity of the organisms that live within them (Ledig, 2002; Young et al., 1993; Keyghobadi, 2007). The mountains of Appalachia in the eastern USA have experienced rapid deforestation over the last four hundred years (Thompson et al., 2013), with forest cover reaching a minimum in in the nineteenth and early twentieth centuries. Over the last century, second-growth forests have replaced once-cleared land as agriculture diminished and forests were protected in the region (e.g. Matlack, 1994; Foster et al., 1998; Bellemare et al., 2002; Gragson and Bolstad, 2006). However, the composition of these eastern forest habitats has been drastically altered, including near extirpation of some species (e.g. of the once dominant American Chestnut) and invasion by alien species (e.g. *Ailanthus alitissima, Lonicera maackii, Microstegium vimineum, and Rosa multiflora*). On top of this, species ranges continue to be altered as temperature and moisture regimes change because of anthropogenic factors.

These changes to Appalachian forests have had profound impacts on the species that comprise them (Matlack, 1994; Stapanian et al., 1998; Flinn and Vellend, 2005). Many have likely experienced severe genetic bottlenecks, followed by explosive increases in abundance (e.g. early or mid-successional forest trees such as beech and maple), while others have recovered slowly or not at all (Vellend, 2004). While the dynamics of forest cover and the major species components of Appalachian forest ecosystems are well studied (i.e. trees), responses of many of the rarer or less conspicuous species occupying Appalachian ecosystems remain relatively poorly studied.

One such species is the leafless, fully mycoheterotrophic, Appalachian orchid *Corallorhiza bentleyi* Freudenstein. This plant was discovered by native orchid enthusiast and author Stan Bentley in the 1990s, and subsequently described as a new species in the genus *Corallorhiza* Gagnebin by Freudenstein (1999). This diminutive, easily overlooked plant is currently known from about a dozen localities across five counties in the eastern US states of Virginia and West Virginia (Freudenstein, 1999; Bentley, 2000; 2004; Virginia Department of Conservation and Recreation, 2009; Barrett and Freudenstein, 2011; Figs. 1, 2). It is found in mid-to-late successional secondary forests, flowering in July-August, often in historically disturbed sites (Bentley, 2000; 2004; Virginia Department of Conservation and Recreation, 2009). Indeed, the type locality for this species is an abandoned logging railroad bed in Monroe County, West Virginia (Freudenstein, 1999). Analysis of morphological and molecular data place *Corallorhiza bentleyi* as a member of the more widespread and morphologically variable *C. striata* Lindl. complex (Freudenstein, 1997; 1999; Barrett and Freudenstein, 2009; 2011). *Corallorhiza bentleyi* shows strong affinities to *Corallorhiza involuta* Greenm., native to southern Mexico, which is similar in stature, characteristics of floral structure (including a putative reproductive mode of autopollination, or selfing), and in DNA sequences. *Corallorhiza bentleyi* differs from *C. involuta* primarily by the size of the callus—two fused lobes at the base of the modified lip petal (labellum)—which is typically larger in *C. bentleyi* (Freudenstein, 1999; Barrett and Freudenstein, 2011). Phylogenetic analysis of plastid (*rbcL, rpl32-trnL*), and nuclear DNA (*ITS, F3H,* and *RPB2*) showed no genetic variation among members of the two species (Barrett and Freudenstein, 2011). Complete plastid genome sequences, however, reveal a pattern of reciprocal monophyly among minimally divergent plastid haplotypes of these two sister species, together which are sister to the remainder of the *C. striata* complex (Barrett et al., 2018).

**Fig. 1.**
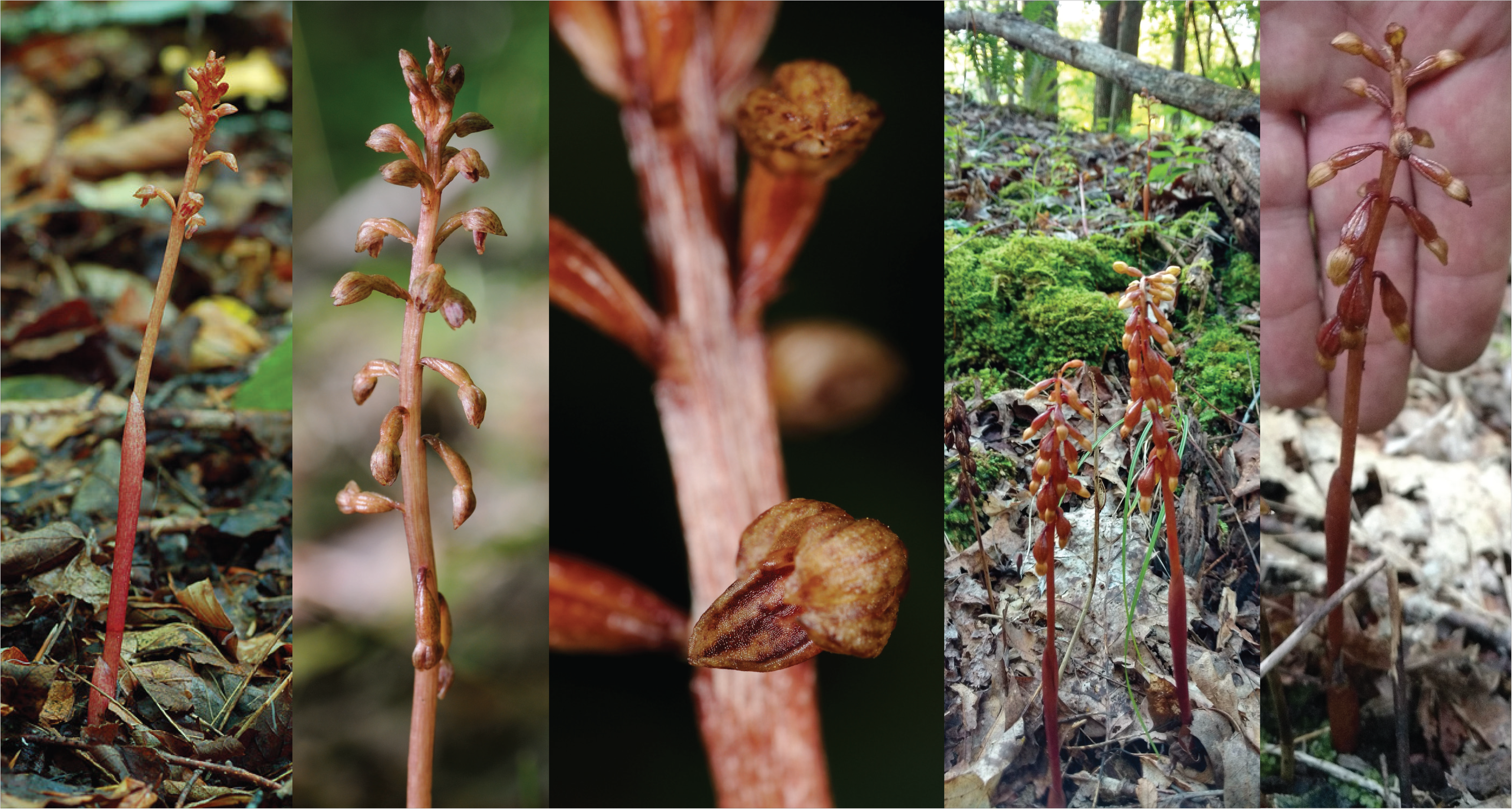
Photos of *Corallorhiza bentleyi*: left three panes, chasmogamous (open-flowered) form; right two panes, cleistogamous form (closed-flowered). Note the early ovary swelling indicative of seed development before the opening of the flowers. Photos: C. Barrett, Z. Bradford)

**Fig. 2.**
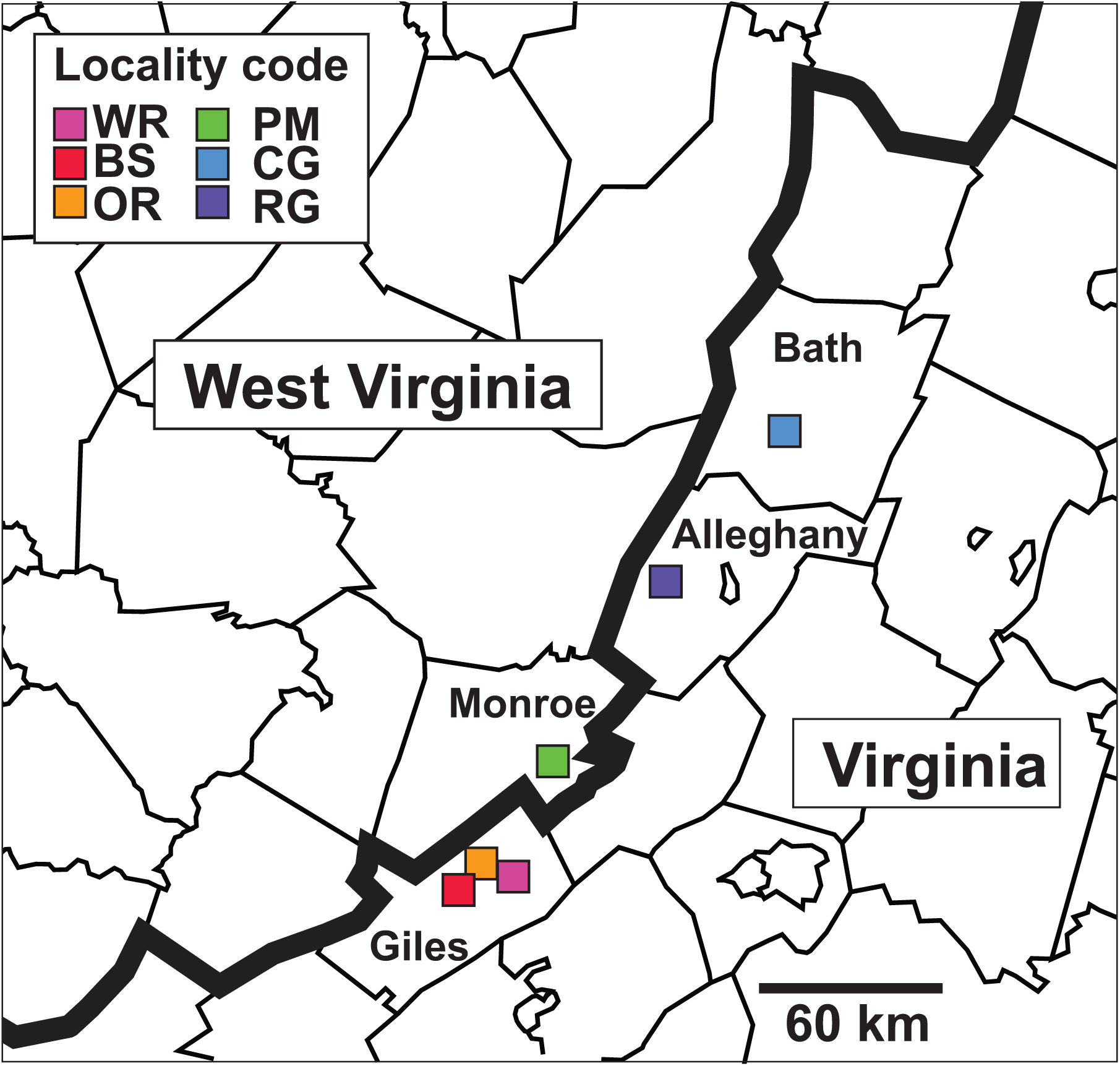
Map of *C. bentleyi* sampling localities by county in Virginia and West Virginia. Legend = 60 km

*Corallorhiza bentleyi* is of high conservation concern. As one of Appalachia’s rarest plants, it is currently classified on the IUCN Red List (https://www.iucnredlist.org/) as ‘vulnerable’ mainly due to its restricted range (a nearly linear, ∼150 km stretch of the Appalachian Mountains in Virginia and West Virginia, USA), low population sizes, and highly fragmented distribution (Bentley, 2004; Goedeke et al., 2015). A survey in 2001 documented ∼400 flowering racemes among all known localities during that time (Bentley 2004, Virginia Department of Conservation and Recreation, 2009). Additionally, *C. bentleyi* shows evidence of obligate self-pollination; the species was originally described as cleistogamous (i.e. flowers remain closed as they self-pollinate) based on specimens from the type locality, but all populations discovered subsequently have open flowers to varying degrees (Bentley, 2004). All flowers in the racemes of *C. bentleyi* are generally self-pollinated by the time they open, as evidenced by swollen ovaries, suggesting early-onset autogamous pollination and seed development. In addition to small population sizes, their autogamous reproductive mode has profound implications for genetic diversity and the ability to adapt to changing environmental conditions. Lastly, this non-photosynthetic orchid completely depends on a rarely collected, poorly understood fungal host for all nutritional needs. Based on sparse sampling of seven *C. bentleyi* individual rhizomes, this species displays specificity towards a single species of ectomycorrhizal fungus in the genus *Tomentella* Pers. ex Pat. (1887), though this small sample is not dense enough to be conclusive (Barrett et al., 2010). Extremely narrow symbiotic specificity thus means that the occurrence and distribution of *C. bentleyi* is completely restricted by that of its host.

Here we take an integrative approach to clarify the conservation genetic status of *C. bentleyi*, using data from floral morphometrics, a conservative and repeatable application of inter-simple sequence repeat (ISSR) variation, and fungal host nuclear internal transcribed spacer region (ITS) sequencing. The goal is to provide a range-wide quantification of genetic, morphological, and symbiotic diversity, and to provide recommendations for the conservation of this ‘vulnerable’ orchid (as currently designated). Specifically, we aim to address several questions regarding the conservation status of this orchid: (1) Are there discernable patterns of morphological differentiation within or among populations of *C. bentleyi*? (2) What is the distribution of genetic diversity within and among the isolated occurrences of *C. bentleyi*? (3) What is the degree of ectomycorrhizal fungal host breadth or specificity across the species range? Lastly, we provide recommendations for the conservation and maintenance of suitable habitat and genetic diversity for this orchid.

## METHODS

### Sampling

We collected samples from 55 individuals of *C. bentleyi* plants across five counties in Virginia and West Virginia, USA (Fig. 1). Small amounts of rhizome tissue (approximately 0.1g of tissue carefully removed from the soil) were collected for 19 individuals, as well as one flower per individual (n = 51 were useable for DNA; n = 47 were useable for morphology), as not to destroy whole plants. Great care was taken to minimize habitat disturbance by trampling and compressing soil in the vicinity of the orchids. For flowers, the perianth was removed from the inferior ovary and fixed in FAA; (formalin 10% v/v), acetic acid 5% v/v, ethanol 50% v/v), and subsequently pickled in 70% ethanol (v/v). One individual flower from each population was kept as a liquid-preserved specimen to represent a voucher at the WVU Herbarium (WVA); whole plants were not collected due to the vulnerability of this species. The type specimen of *C. bentleyi* is housed at the Oakes Ames Orchid Herbarium (AMES) at Harvard University Herbaria, corresponding to the ‘PM’ locality from the current study in Monroe Co., WV (Freudenstein, 1999).

### Morphology

We measured 14 floral characters for 47 individuals of *C. bentleyi* with a Fisherbrand™ Digital Stereo Microscope (Fischer Scientific, Waltham, Massachusetts, USA) calibrated to a 2,000 μm slide, and analyzed each flower digitally using the Image J software package (Abràmoff et al., 2004). We measured: Column length (the gynostemium, or the fused stamen and gynoecium); column apical width; column basal width; column width at narrowest point; dorsal sepal length; dorsal sepal width; lateral sepal length; lateral sepal width; lateral petal length; lateral petal width; labellum length (i.e. the lip, a modified, pendulous petal); labellum width at widest point; callus length (two fused lobes at the base of the labellum); and callus width at widest point. Morphological analyses were conducted via Principal Components Analysis (PCA) in PAST v.3 (Hammer et al., 2001), using a correlation matrix to standardize the scale of measurements among characters, and nonparametric multivariate analysis of variance (NPMANOVA, n = 9,999 permutations on a log_10_-transformed matrix of Euclidean distances) to test for distinctness in morphology among sampling localities. Pairwise comparisons in the NPMANOVA analysis were Bonferroni-corrected.

### Fungal host DNA

We extracted combined plant and fungal DNA from rhizome tissues of 19 individuals (see above). We used primers ITS1F and ITS4 (White, *et al.,* 1990; Gardes and Bruns, 1993) to selectively amplify fungal DNA from the Internal Transcribed Spacer region (ITS). The PCR settings for all samples followed: initial template denaturation at 95°C for 5 minutes; 30 cycles of denaturation at 95°C for 30 seconds, primer annealing at 55°C for 30 seconds, and primer extension at 72°C for 1 minute; and final primer extension at 72°C for 5 minutes. Excess PCR reagents were removed using an AxyPrep Fragment Select magnetic beads (Axygen, Big Flats, New York, USA) and two washes with 80% ethanol (v/v). Cycle sequencing reactions were conducted as in Barrett et al. (2010) using the BigDye v.3.1 cycle sequencing kit (Applied Biosystems, Foster City, California, USA). We removed excess dyes and primers using Sephadex G-50 fine medium and centrifugation through a filter plate (Phenix Research, Accident, Maryland, USA). Cleaned reactions were then run at the WVU Genomics Core Facility on an Applied Biosystems 3130xl Genetic Analyzer. Electropherograms were assembled *de novo* under default settings and edited in Geneious v.10 (www.geneious.com). Cloning of PCR products was not necessary, as *Corallorhiza* individuals nearly always associate with a single fungal ITS type (see Taylor et al., 2004; Barrett et al., 2010; Freudenstein and Barrett, 2014). Fungal ITS sequences from the 19 *C. bentleyi* individuals were subjected to BLAST sequence analysis via the NCBI database (https://www.ncbi.nlm.nih.gov/) for fungal host identification. We used default BLAST parameters (MEGABLAST), but excluded environmental samples, as these are often of uncertain taxonomic affinity or are poorly annotated. We chose a minimum e-value of 10^-5^, and chose the best hit as the sequence from NCBI with the lowest e-value and maximum BLAST score, requiring a full species epithet in the annotation (i.e. species names of “sp.” were not considered).

Sequences were then aligned in MUSCLE (Edgar, 2004) with default parameters. We then re-aligned the 19 *C. bentleyi* sequences with along with 101 fungal ITS sequences from the *C. striata* complex (Barrett and Freudenstein, 2010), and including 1,433 reference sequences from NCBI GenBank (https://www.ncbi.nlm.nih.gov/genbank), representing all currently sequenced species of the genus *Tomentella* using MAFFT (using the ‘auto’ algorithm, with the gap opening parameters set to 2.5 and he offset parameter set to 0.5). We excluded unvouchered, environmental samples of uncertain taxonomic affinity. Sites with >95% gaps were masked, corresponding to poorly aligned regions and homopolymer runs, resulting in a final aligned matrix of 807 positions. The close relative *Thelephora terrestris* (GenBank accession number UDB000216) was chosen as the outgroup (see Online Resource 1 for final alignment). We used the GTR-GAMMA model of sequence evolution to build a phylogenetic tree in RaxML v.8 (Stamatakis 2008), with 1,000 standard bootstrap replicates, repeating the tree search ten times from random starting seeds for the final ML estimate of topology and branch lengths.

### Genetic diversity

We used a modified CTAB method to extract genomic DNA from fresh ovary (shoot) tissue from 51 individuals, with RNAse A treatment (Doyle and Doyle, 1987). We removed ∼0.1g fresh or frozen tissue from the outer portion of ovary tissue, being careful not to sample seed tissues. Samples were stored at −80C until further use. Initial attempts to identify polymorphic microsatellite markers, designed using Illumina short-read data from a previous study (Barrett et al., 2018) were unsuccessful, with all microsatellite regions being monomorphic. Thus, we took an alternative approach using inter-simple sequence repeats (ISSR). 25 ISSR primers from the University of British Columbia ISSR Set (Goh *et al.,* 1999) were initially amplified in single-primer reactions using Polymerase Chain Reaction (PCR) on a panel of six *C. bentleyi* individuals to identify variable banding patterns among individuals. The dominant nature of ISSR markers is less ideal than codominant markers such as microsatellites, especially for estimating parameters such as observed heterozygosity and linkage disequilibrium. However, dominant ISSR markers are still highly repeatable, and useful for determining levels of genetic diversity within and among sampling localities in the absence of suitable or feasible codominant markers.

PCR settings for all ISSR samples were conducted following: 95°C for 4 min; 35 cycles of denaturation at 94°C for 30 seconds; primer annealing at 50°C for 45 seconds; and primer extension at 72°C for 2 minutes. The program was held at 72°C for an additional 10 minutes of primer elongation, then stabilized at 4°C. Two ISSR primers were ultimately chosen and amplified for all *C. bentleyi* samples, based on their initial levels of variation and ease of scoring: UBC-868 (GAA)_6_ and UBC-873 (GACA)_4,_ using the same PCR conditions. Samples from 47 individuals for both primers were amplified and run on polyacrylamide or agarose gels to observe ISSR banding patterns for each individual. In order to validate the exact length and intensity of each band, we took a conservative approach by rendering digital gel images for the same 47 individuals using a 2100 Bioanalyzer (Agilent, Santa Clara, California, USA). As a result, we were able to confirm banding patterns found in agarose and polyacrylamide gels, as well as to verify bands of lower intensity across samples. All reactions were run in duplicate to confirm repeatability of band presence/absence (see Online Resource 2A). Bands were scored visually from gels and Bioanalyzer traces (e.g. Online Resource 2B,) and then were used to populate a binary presence/absence matrix. We consider this approach to be conservative and consistent in terms of both repeatability and accuracy. Inclusion of bioanalyzer traces allowed us to compare band sizes relative to a known standard with high precision (i.e. down to a few base pairs for each peak), reducing uncertainty in homology assessment of bands.

ISSR bands were scored as present (1) or absent (0) at each base pair along the digital gel and compared with respective agarose and polyacrylamide gels. Bands with a total frequency of >95% or < 5% were excluded. We used GenAlEx v.6.5 (Peakall and Smouse, 2006; 2012) to analyze the binary matrix created from presence/absence scoring (Online Resource 3). We conducted Analysis of Molecular Variation (AMOVA) using the binary data option for dominant markers. We conducted 9,999 permutations to assess statistical significance of observed values at three levels: (1) regional, i.e. northern vs. southern localities (i.e. Bath and Alleghany Counties, Virginia vs. all other localities; Fig. 2); (2) among sampling localities; and (3) among individuals within sampling localities. We also used GenAlEx to conduct Mantel tests for correlation between geographic and observed genetic distances between sampling localities (using Φ-statistics, the haploid or dominant-marker analog to F-statistics). Mantel tests were run for 9,999 permutations using linear genetic and geographic distance matrices, the latter of which was derived from decimal-formatted coordinates. GPS coordinates are protected due to the IUCN ‘vulnerable’ status of this orchid. We further analyzed ISSR genetic variation among *C. bentleyi* individuals via Principal Coordinates Analysis (PCoA) and NPMANOVA in PAST v.3 (Hammer et al., 2001), based on a binary matrix with Jaccard distance transformation. Significance of NPMANOVA was calculated via 9,999 permutations, and pairwise differences were calculated with Bonferroni correction. We tested whether each sampling locality harbored significantly different levels of genetic variation among sampling localities by subjecting the PCoA scores among individuals for each locality to Levene’s test for homogeneity of variances in PAST v.3. We conducted Levene’s test for individual sample scores on PCoA axes 1 and 2, grouped by sampling locality. Lastly, we constructed a dendrogram relating overall genetic similarity among different sampling localities using Nei’s unbiased genetic diversity (Nei, 1978) in POPGENE v.1.32 (Yeh et al., 1999).

All data for ISSR banding and morphology are deposited in Dryad (link, XXXXXX).

## RESULTS

### Morphology

Principal Components Analysis of morphometric characters identified distinct groupings among some sampling localities (Fig. 3). PC1 accounted for 66.9% of variability and PC2 accounted for 10.3% in the floral data. Individuals from the northernmost (CG, Bath, VA) and southernmost localities (OR and BS, Giles, VA; Fig. 3A) clustered closely, with little overlap otherwise between populations. Populations within Alleghany County (RG) showed the most within-locality variation along PC1, while plants from the WR locality (Giles Co., VA) showed the most variation among PC2; individuals from Monroe, WV (PM) showed relatively little variation. All floral characters were somewhat positively associated with PC1, whereas column width (apex), callus length and width, and labellum length and width contributed positively to PC2 (Fig. 3B.). Column length and width characters contributed negatively to PC2, whereas all the remaining characters showed little variation with respect to PC2.

**Fig. 3.**
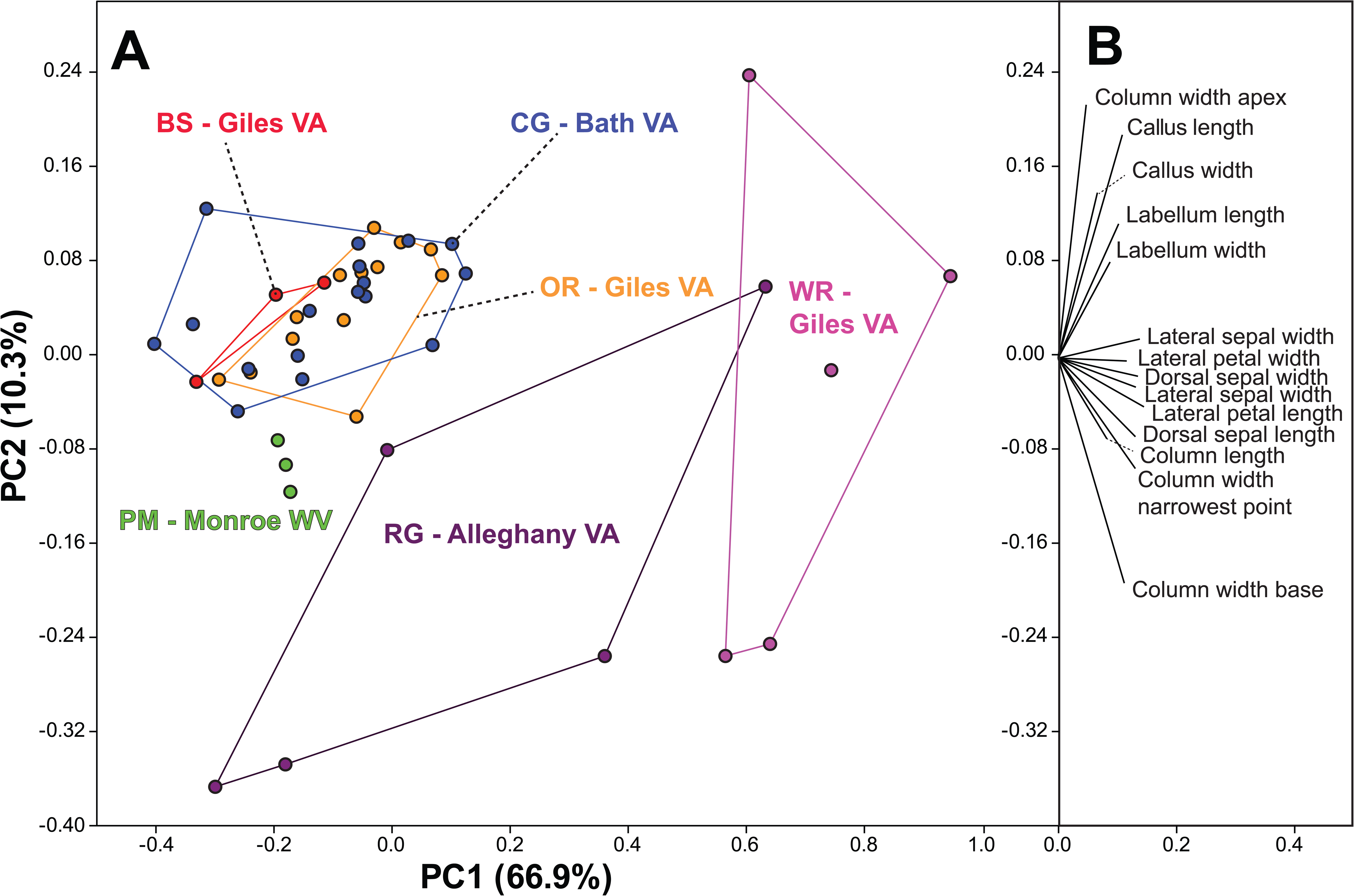
**A** Principal components analysis (PCA) scatterplot of 14 quantitative floral characters, transformed via a correlation matrix, showing principal components 1 & 2. Lines around points are convex hulls corresponding to each color-coded collection locality. **B** Biplot of each character’s correlation with PC 1 and 2

### Fungal hosts

All nineteen rhizome samples were successfully amplified and sequenced with an average ITS length of 679 base pairs (GenBank accession numbers XXXXXX-XXXXXX). NCBI BLAST searches confirmed the 19 sequences were from the ITS region, and each most closely matched one of five fully annotated *Tomentella* species from NCBI GenBank (Table 1). Ten of the nineteen samples most closely matched a single accession identified as *T. fuscocinerea* (Pers.) Donk from Iran (accession GU214810, 93.50-97.00% identity); four matched most closely matched an ectomycorrhizal associate of *Pinus koraiensis* Siebold & Zucc. from Russia (identified as *T. lateritia* Pat., accession KP783474, 93.05-98.00% identity); four matched a soil isolate identified as *T. fuscocinerea* from Alaska, USA (KY322557, 93.53-97.41% identity); and one matched an associate of *Crataegus monogyna* Jacq. identified as *T. fuscocinerea* from Portugal (accession FN594893; 97.56% identity).

**Table 1.** BLAST Results for *C. bentleyi* fungal ITS sequences. Results displayed are those of the closest named species matching each *C. bentleyi* sequence. Environmental samples were excluded from the search, which was conducted under default parameters via the NCBI web interface. ‘Acc.’ = collection number for each accession sequenced; ‘Source’ = information available from GenBank on the host and locality collected for each match, ‘Query Cover’ = the percent coverage of the ITS sequence; ‘%ID’ = percent identity to closest match; ‘GB’ = GenBank accession number of the closest matching accession with a species-level identification.

| Acc. | Locality,<br>county | Description | Source | Query<br>Cover | % ID | GB |
| --- | --- | --- | --- | --- | --- | --- |
| **63aR** | BS, Giles,<br>VA | *Tomentella fuscocinerea*<br>isolate S29.1 | Soil, Alaska, USA | 98% | 93.55% | KY322557.1 |
| **65aR** | BS, Giles,<br>VA | *Tomentella fuscocinerea*<br>isolate S29.1 | Soil, Alaska, USA | 100% | 93.53% | KY322557.1 |
| **G5R** | OR, Giles,<br>VA | *Tomentella lateritia*<br>isolate 403 | *Pinus koraiensis*,<br>Russia | 98% | 93.19% | KP783474.1 |
| **B6R** | CG, Bath,<br>VA | *Tomentella fuscocinerea*<br>isolate S29.1 | Soil, Alaska, USA | 95% | 93.65% | KY322557.1 |
| **68aR** | BS, Giles,<br>VA | *Tomentella lateritia*<br>isolate 403 | *Pinus koraiensis*,<br>Russia | 99% | 93.05% | KP783474.1 |
| **64R** | BS, Giles,<br>VA | *Tomentella fuscocinerea*<br>voucher TU108229 | Iran | 96% | 93.52% | GU214810.1 |
| **67R** | BS, Giles,<br>VA | *Tomentella lateritia*<br>isolate 403 | *Pinus koraiensis*,<br>Russia | 98% | 93.32% | KP783474.1 |
| **423bR** | WR, Giles,<br>VA | *Tomentella fuscocinerea*<br>voucher TU108229 | Iran | 97% | 96.46% | GU214810.1 |
| **423cR** | WR, Giles,<br>VA | *Tomentella fuscocinerea*<br>voucher TU108229 | Iran | 97% | 96.46% | GU214810.1 |
| **B19R** | CG, Bath,<br>VA | *Tomentella fuscocinerea*<br>voucher TU108229 | Iran | 96% | 96.76% | GU214810.1 |
| **B4R** | CG, Bath,<br>VA | *Tomentella fuscocinerea*<br>voucher LISU:178262 | *Crataegus*<br>*monogyna*,<br>Portugal | 94% | 97.56% | FN594893.1 |
| **426cR** | RG,<br>Alleghany,<br>VA | *Tomentella fuscocinerea*<br>voucher TU108229 | Iran | 97% | 94.78% | GU214810.1 |
| **424bR** | PM,<br>Monroe,<br>WV | *Tomentella fuscocinerea*<br>voucher TU108229 | Iran | 97% | 96.13% | GU214810.1 |
| **424aR** | PM,<br>Monroe,<br>WV | *Tomentella fuscocinerea*<br>voucher TU108229 | Iran | 97% | 96.30% | GU214810.1 |
| **423aR** | WR, Giles,<br>VA | *Tomentella fuscocinerea*<br>voucher TU108229 | Iran | 98% | 96.46% | GU214810.1 |
| **424cR** | PM,<br>Monroe,<br>WV | *Tomentella fuscocinerea*<br>isolate S29.1 | Soil, Alaska, USA | 96% | 97.41% | KY322557.1 |
| **426bR** | RG,<br>Alleghany,<br>VA | *Tomentella fuscocinerea*<br>voucher TU108229 | Iran | 97% | 94.90% | GU214810.1 |
| **426aR** | RG,<br>Alleghany,<br>VA | *Tomentella fuscocinerea*<br>voucher TU108229 | Iran | 97% | 94.73% | GU214810.1 |
| **G18R** | OR, Giles,<br>VA | *Tomentella lateritia*<br>isolate 403 | *Pinus koraiensis*,<br>Russia | 98% | 93.19% | KP783474.1 |

Phylogenetic analysis of fungal ITS sequences among 19 *C. bentleyi* associates and >1,400 reference sequences revealed that the sampled individuals clustered most closely with sequences of *Tomentella fuscocinerea,* with some accessions of *Tomentella patagonia* Kuhar & Rajchenb. and other, unidentified *Tomentella* spp. (Fig. 4A). Furthermore, all associates of *C. bentleyi* and the more geographically widespread *C. striata* complex formed a clade with all accessions of the ‘*Tomentella fuscocinerea-patagonica’* complex (Fig. 4A; bootstrap = 0.89), within which all *C. bentleyi* fungal associates grouped in one of four distinct clades (Fig. 4B). The first comprised five accessions from three localities (BS, GC, and OR; bootstrap = 1.0) closely related to two *Tomentella* spp. accessions from Svalbard, Norway and other accessions of *C. striata* from Arizona, New Mexico, and Michigan, USA (Fig. 4B). The second clade consisted of five accessions from three localities (CG, PM, and WR; bootstrap = 0.74), closely related to *Tomentella* accessions (including two accessions identified as *T. fuscocinerea*) from Iran, Hungary, Montenegro, and Minnesota, USA, as well as from *C. striata* associates from Wyoming and California, USA. A third, relatively weakly supported clade consisted of eight *C. bentleyi* associates from three localities (BS, PM, and RG, bootstrap = 0.62), sister to a large clade of *C. striata* accessions sampled from across North America (bootstrap = 1.0). Lastly, a single accession (B4R, from locality CG) clustered with three *T. fuscocinerea* isolates and several accessions from other members of the *C. striata* complex (bootstrap value = 1.00).

**Fig. 4.**
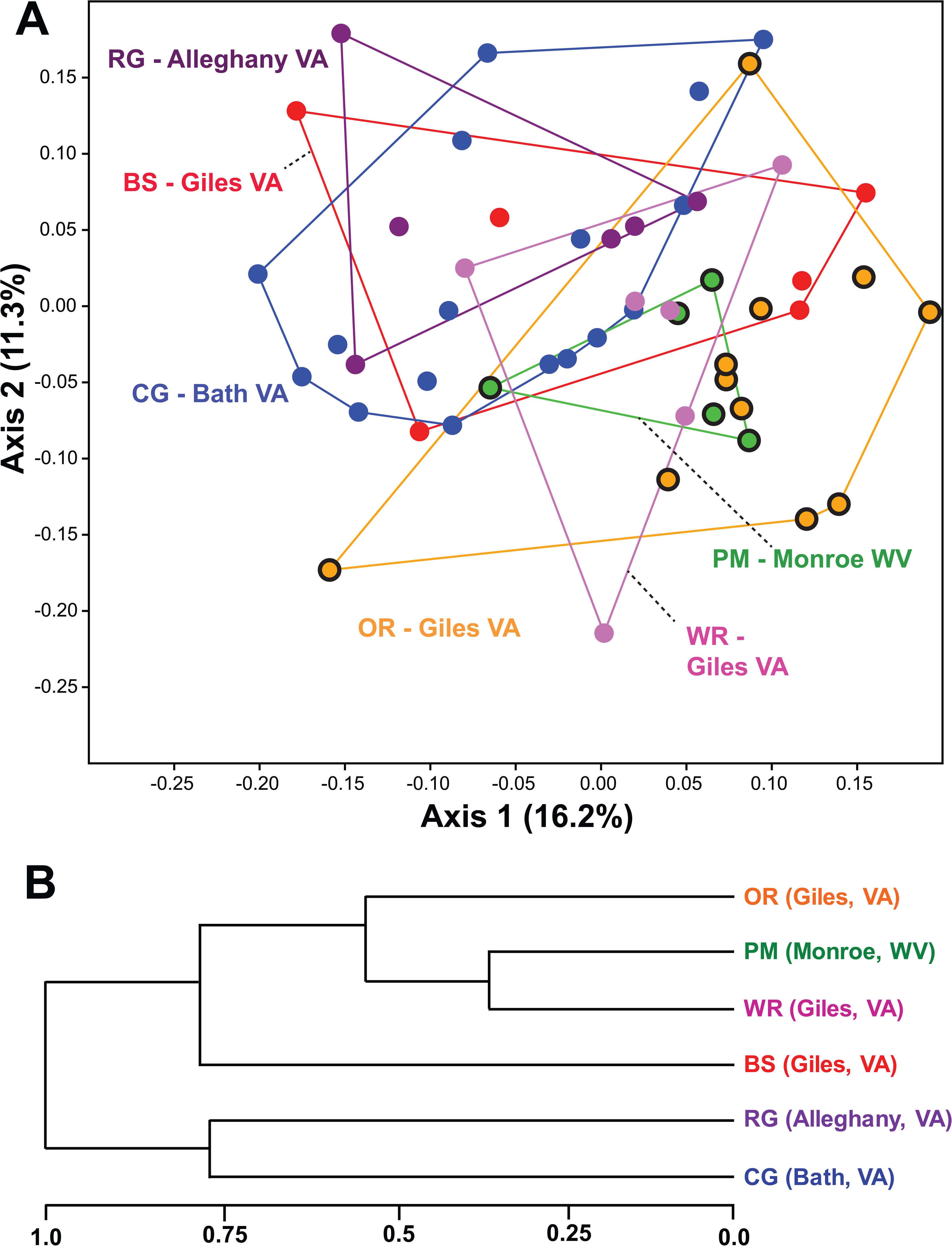
Fungal ITS Maximum Likelihood tree of *C. bentleyi* fungal associates. **A**. Phylogram of 1,452 sequences of Thelephoraceae from GenBank in addition to *C. bentleyi* and other associates of the *C. striata* complex. Scale bar = 0.2 substitutions/site. **B.** Phylogenetic relationships of *C. bentleyi* associates. Numbers above branches = ML bootstrap percentages. Arrows correspond to the four clades in which *C. bentleyi* associates were placed

### Genetic diversity

Of the 25 ISSR primers initially screened (Pharmawati *et al.,* 2005), primers 873 and 868 were the most informative and showed variable bands for *C. bentleyi* individuals. Polyacrylamide gel images of individuals amplified with primer 873 showed a range of 10-30 distinguishable bands, with bands consistently present at 525bp, 900bp, and 970bp. Agarose gel images of individuals amplified with primer 868 showed a range of 9-20 distinguishable bands, with bands consistently present at 275bp, 600bp, and 675bp. We scored 48 total loci for primer 873, and 41 total loci for primer 868.

Overall metrics of genetic diversity for ISSR loci are presented in Table 2. The mean percentage of polymorphic loci among sampling localities was 80.89% (S.E. = 4.23%), with a range of 68% in PM (Monroe Co., WV) to 97.3% in CG (Bath Co., VA). Overall, locality CG in Bath, VA had the highest measures of genetic diversity (percentage of polymorphic loci, Shannon’s Diversity Index, haploid binary genetic diversity, and unbiased haploid binary genetic diversity; Table 2). Locality PM (the type locality in Monroe Co., WV) had the lowest diversity; the exception was that locality OR had lower unbiased genetic diversity, when corrected for sample size differences among localities. We found no evidence of identical genomic clones among individuals within or between localities for the 75 polymorphic loci scored.

**Table 2.** Summary statistics across 75 polymorphic loci for six *C. bentleyi* localities. ‘N’ = number of individuals sampled per locality; ‘%p’ = percentage of polymorphic loci; ‘I’ = Shannon’s Diversity Index; ‘h’ = haploid binary genetic diversity (h = 1 – Σp_i_^2^ where ‘p’ indicates band presence at the ‘i^th^’ locus); ‘uh’ = unbiased haploid diversity (uh = h*(N/(N – 1)).

| Locality |  | N | %p | I | h | uh |
| --- | --- | --- | --- | --- | --- | --- |
| OR (Giles, VA) | Mean | 11 | 81.33 | 0.420 | 0.279 | 0.306 |
|  | SE |  |  | 0.028 | 0.021 | 0.023 |
| PM (Monroe, WV) | Mean | 5 | 68.00 | 0.379 | 0.254 | 0.317 |
|  | SE |  |  | 0.031 | 0.021 | 0.027 |
| BS (Giles, VA) | Mean | 7 | 85.33 | 0.476 | 0.320 | 0.373 |
|  | SE |  |  | 0.026 | 0.019 | 0.022 |
| WR (Giles, VA) | Mean | 6 | 81.33 | 0.451 | 0.302 | 0.363 |
|  | SE |  |  | 0.027 | 0.020 | 0.023 |
| RG (Alleghany, VA) | Mean | 5 | 72.00 | 0.427 | 0.292 | 0.365 |
|  | SE |  |  | 0.032 | 0.023 | 0.028 |
| CG (Bath, VA) | Mean | 17 | 97.33 | 0.560 | 0.381 | 0.405 |
|  | SE |  |  | 0.017 | 0.014 | 0.015 |
| Total | Mean | 8.5 | 80.89 | 0.452 | 0.305 | 0.355 |
|  | SE | 0.2 | 4.23 | 0.011 | 0.008 | 0.010 |

Principal coordinates axis 1 explained 16.2% of the Jaccard-transformed ISSR band presence/absence variation, while axis 2 explained 11.3% (Fig. 5A). Overall there was no evidence for clear separation among localities in multidimensional space based on ISSR loci. Localities PM and WR, however, occupy smaller areas of multidimensional space relative to those from individual localities, revealing lower overall genetic diversity associated with these two axes. Levene’s Test for homogeneity of variances was marginally non-significant for individual scores on PCoA axis 1 (p = 0.063) but was significant for scores on axis 2 (p = 0.028), a proxy for differences in levels of genetic diversity. Nonparametric MANOVA indicated significant differentiation among localities overall (F=1.996; p = 0.0001). Pairwise, Bonferroni-corrected NPMANOVA (Table 3) differentiated locality OR (Giles, VA) from CG (Bath, VA) and RG (Alleghany, VA). A dendrogram based on Nei’s unbiased genetic distance (Fig. 5B) placed PM (Monroe, WV) and WR (Giles, VA) as most genetically similar, followed by OR and BS (Giles, VA, respectively); together these represent the southernmost localities, while the two northernmost localities RG (Alleghany, VA) and CG (Bath, VA) group together and are more genetically dissimilar to the southern localities (Fig. 5B).

**Fig. 5.**
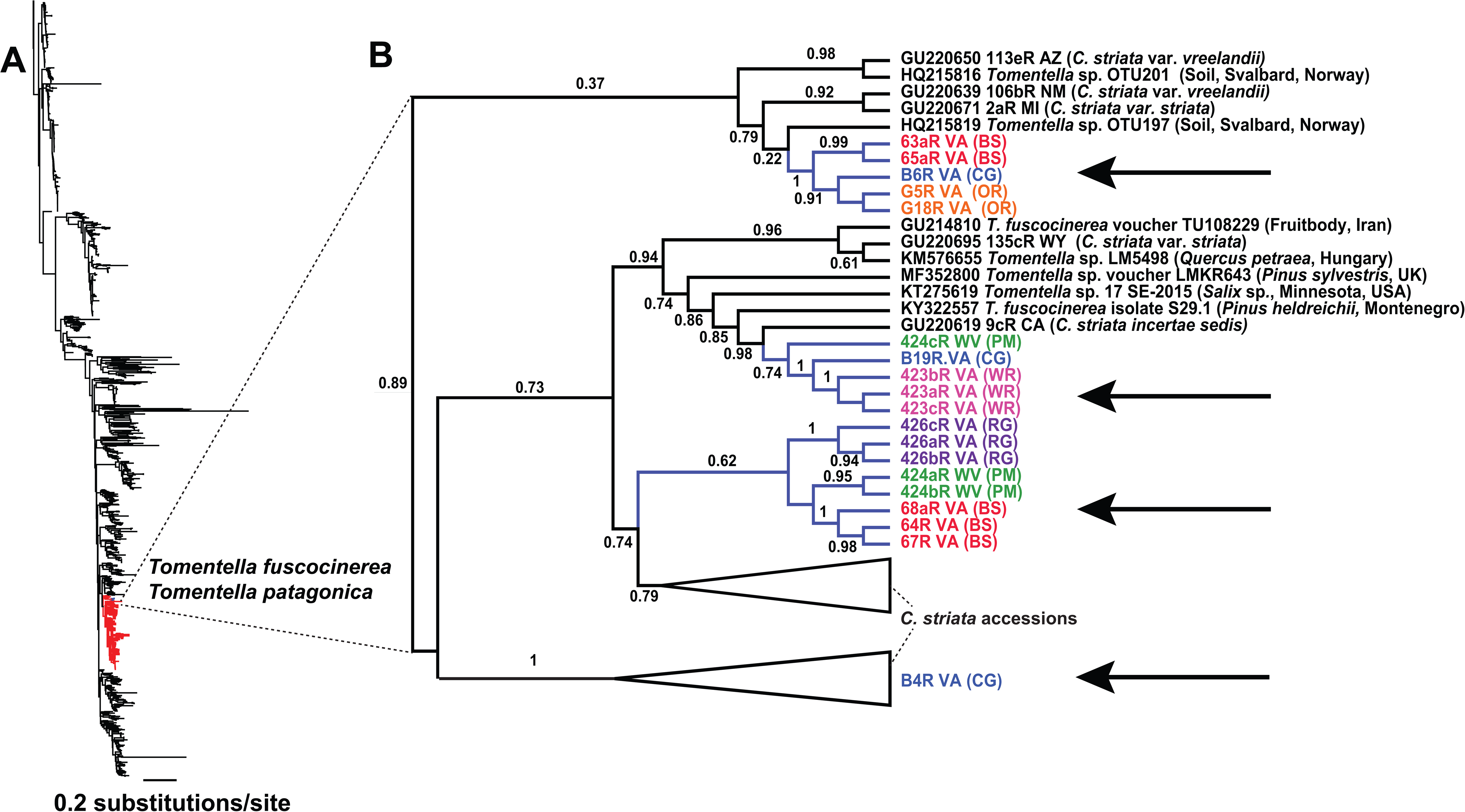
**A.** Principal Coordinates analysis of ISSR band presence/absence, coded as binary characters, and transformed via Jaccard distance. Lines around points are convex hulls corresponding to each color-coded collection locality. **B.** Dendrogram of Nei’s unbiased genetic distance among sampling localities.

**Table 3.**
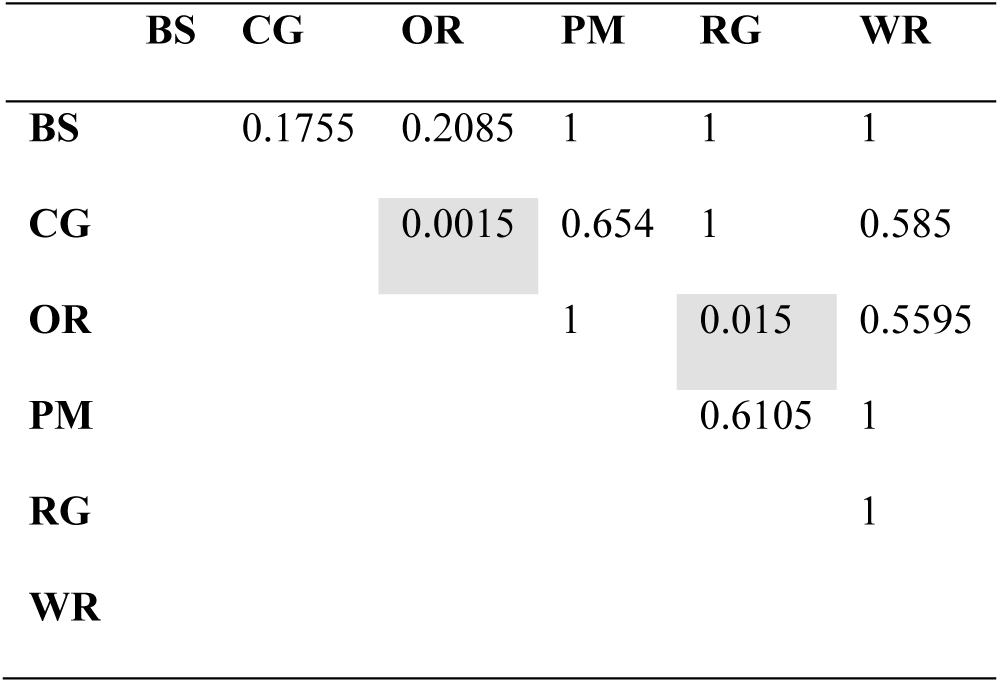
Pairwise, Bonferroni-corrected p-values based on 9,999 permutations (overall total sum of squares: 9.234; within-group sum of squares: 7.558; F-value: 1.996; P-value: 0.0001). Gray shading = significant P-values (P < 0.05).

Results from analysis of molecular variance (AMOVA) revealed that 89% of total molecular variance among ISSR loci occurs within sampling localities, six percent occurs between northern and southern regions (northern populations = CG and RG, southern = WR, BS, OR, and PM), and the remaining five percent occurs among localities (Table 4). The between-regional Phi_RT_ estimate was 0.059 (p = 0.001), among populations Phi_PR_ within regions was 0.052 (p = 0.006), and within sampling localities Phi_PT_ was 0.108 (p < 0.0001). Overall observed Phi_PT_ fell outside the 95% confidence interval of 9,999 randomized permutations. Plants from locality RG (Alleghany, VA) had the highest pairwise Phi_PT_ values compared to all other localities, indicating it is the most genetically distinct from all others (Table 5). Mantel analysis of overall genetic vs. geographic distances among localities was significant (R = 0.231, p < 0.0001, 9,999 permutations), indicating that individuals were genetically more similar to others from localities closer in geographic proximity along a slight north-to-south gradient.

**Table 4.**
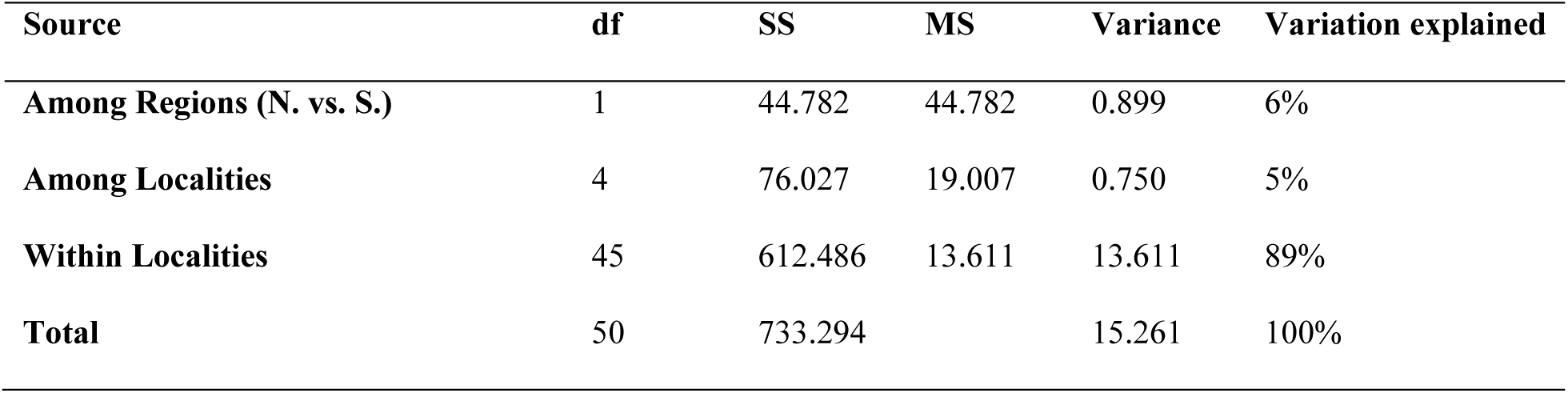
Analysis of Molecular Variance (AMOVA) from 75 polymorphic ISSR loci. ‘df’ = degrees of freedom (n-1); ‘’SS’ = sum of squares; ‘MS’ = mean of squares; ‘Variation explained’ = the total proportion of variation explained by each hierarchical component. ‘N. vs. S.’ defines localities in the northern (Bath Co., VA, Alleghany Co., VA) and southern (Giles Co., VA, Monroe Co., WV) portions of the range. The northernmost population was unsampled (Pocahontas Co., WV).

**Table 5.** Pairwise Phi_PT_ values between sampling localities. Below diagonal = Phi_PT_ values; above diagonal = significance of Phi_PT_ values after Bonferroni correction. Gray shading indicates significance at the p = 0.05 level.

|  | OR | PM | BS | WR | RG | CG |
| --- | --- | --- | --- | --- | --- | --- |
| OR |  | 0.104 | 0.004 | 0.023 | 0.001 | 0.000 |
| PM | 0.040 |  | 0.047 | 0.456 | 0.097 | 0.063 |
| BS | 0.103 | 0.081 |  | 0.147 | 0.031 | 0.006 |
| WR | 0.061 | 0.000 | 0.038 |  | 0.106 | 0.071 |
| RG | 0.209 | 0.096 | 0.105 | 0.072 |  | 0.090 |
| CG | 0.133 | 0.058 | 0.093 | 0.051 | 0.054 |  |

AMOVA of individual ISSR loci revealed seven loci that accounted for maximal genetic differentiation at the locality and regional levels (Table 6). The presence or absence of these ISSR loci was either unique to northern or southern regions, unique to certain sampling localities, or accounted for diversity within a locality. The numbering of loci here is based on order of base pair length for the 75 total ISSR loci analyzed via Bioanalyzer. Locus 1 occurs almost exclusively in the southern region (OR, BS, and WR; Table 6). Locus 7 is present primarily in the northern region, almost exclusively in locality CG. Locus 8 is present in all WR and RG individuals (Giles and Alleghany, VA, respectively), as well as in at least one individual of every other locality. Locus 15 is most prevalent in the northernmost locality of CG (Bath, VA) and less so in southern localities. Locus 15 is absent from RG (Alleghany, VA), but is variably present at all other localities. Locus 34 is found primarily in southern localities and is present in every individual in the PM population. Loci 41 and loci 49 were most prevalent in CG (Bath, VA) and RG (Alleghany, VA), the two northernmost localities, and showed relatively lower frequency in southern localities.

**Table 6.** Phi-statistics from AMOVA of 75 polymorphic ISSR loci. Phi_RT_ = among regions (North vs. South); Phi_PR_ = among localities within regions; Phi_PT_ = among individuals within localities. ‘Total’ includes all 75 loci, followed by all individual loci with significant values for at least one Phi-statistic. One, two, or three asterisks indicate significance at the p = 0.05, 0.01, and 0.001 levels, respectively.

|  | **Total** | **Locus 1** | **Locus 7** | **Locus 8** | **Locus 15** | **Locus 34** | **Locus 41** | **Locus 49** |
| --- | --- | --- | --- | --- | --- | --- | --- | --- |
| **$\Phi_{RT}$** | 0.059*** | 0.477*** | 0.233** | -0.340 | 0.085 | 0.353*** | 0.206* | 0.389** |
| **$\Phi_{PR}$** | 0.052** | 0.816*** | 0.375** | 0.375** | 0.216* | 0.392*** | 0.171* | 0.307** |
| **$\Phi_{PT}$** | 0.108*** | 0.648*** | 0.185* | 0.534*** | 0.143 | 0.061 | -0.043 | -0.134 |

## DISCUSSION

### Morphology

There is evidence for morphological differentiation among sampling localities: RG and WR from Alleghany and Giles, VA displayed the most variation among PCs 1 and 2 (Fig. 3A). Locality PM (Monroe, WV), on the other hand, had the lowest degree of morphological variation overall; interestingly, localities from which more samples were collected did not yield higher levels of variation (e.g. CG from Bath, VA, and OR from Giles, VA had the highest sample sizes but relatively low levels morphological variation). Locality PM, the type locality of *C. bentleyi*, is the only known population in which all individuals are completely cleistogamous (i.e. flowers never open). It is unsurprising then that morphological variation would be the lowest here, as this condition completely prevents any chance of outcrossing.

Field-collected material that is not subjected to common garden or reciprocal transplant experimentation likely reflects spatiotemporal variation in environmental factors such as rainfall, temperature, soil properties, or factors affecting photosynthate output of the trees that nourish their fungal hosts. Common garden experiments are extremely difficult if not impossible with mycoheterotrophs such as *C. bentleyi* due to their strict tripartite symbiotic requirements, slow growth rates (e.g. see Rasmussen, 1995), and potential biases associated with plant-fungal-autotroph compatibility/fitness issues (in tree seedling microcosm experiments, e.g. McKendrick et al., 2000). While characteristics such as plant height/biomass, the number of flowers per raceme, seed production (i.e. fecundity), and other factors might result from habitat quality *in situ*, it is more difficult to imagine how or why such biotic and abiotic factors might influence characters such as floral length/width dimensions, which would intuitively seem to be more under genetic control. One possibility would be a simple allometric scaling effect: sizes of floral parts may be correlated with total plant size, allowing an explanation based on environmental variables to be taken into consideration (Niklas, 1994; 2004). Unfortunately, due to the vulnerable status of this orchid, metrics such as biomass would require destructive sampling which is contra to the conservation purposes of this study.

The other possibility is that variation in floral characters is at least somewhat heritable, and possibly adaptive. One obvious reason for this would be regional or habitat-associated variation in pollinators (e.g. Price et al., 2005; Reverte et al., 2019). Such variation is observed in many taxa, even in other species complexes within *Corallorhiza*; e.g. cleistogamous and chasmogamous forms of *C. odontorhiza*, and nearly clinal variation of floral size and display in *C. striata* vars. *striata* and *vreelandii* across north America, the latter possibly associated with variation in the degree of selfing among populations (Freudenstein, 1997; Barrett and Freudenstein, 2009; 2011; Freudenstein and Barrett, 2014). Some members of the *C. striata* complex—to which *C. bentleyi* belongs—are known to be pollinated by male ichneumonid wasps (Freudenstein, 1997; Barrett and Freudenstein, 2009), which appear to play more of an important role in the predominantly outcrossing populations of *C. striata* Lindl. var. *striata* in the northern USA and Canada. The latter have large, open, showy flowers and do not set seed with the frequency of other members of this complex (e.g. *C. striata* var. *vreelandii* (Rydb.) L.O. Williams, *C. involuta*, and *C. bentleyi*; Freudenstein, 1997; Barrett and Freudenstein, 2014). There is anecdotal evidence that male ichneumonid wasps may visit open flowers of *C. bentleyi* based on a single photograph (taken by Charles Garratt, Warm Springs, VA, USA), but there are no recorded pollinators of *C. bentleyi* to date despite several hours of observations of the plants in flower (C. Barrett and J. Freudenstein, personal observation). Thus, it is unclear if or how floral variation among sampling localities, or even genotypes within localities might be driven by pollinator specificity, given the observed propensity of this species to self-pollinate.

### Fungal hosts

As with other members of the *C. striata* complex, *C. bentleyi* associates with an extremely narrow clade of ectomycorrhizal fungi in the genus *Tomentella*, namely *T. fuscocinerea*. This pattern holds across six sampling localities spanning the geographic distribution of the species. Little is known about *T. fuscocineara*, including its preferred habitat, soil requirements, or fidelity to specific photobiont hosts. It has been sporadically collected on several continents, implying a global distribution. Yet, it is rarely collected in fruit, even in environmental and soil samples, suggesting low abundance and an extremely patchy distribution; indeed, only 14 records of *T. fuscocinerea* exist in NCBI GenBank, with 131 putative records in the Global Biodiversity Information Facility (GBIF; https://www.gbif.org/). Though our sample of only 19 *C. bentleyi* rhizomes is not large (mainly due to the physical difficulty of obtaining rhizome material without destroying plants), we were able to infer that *C. bentleyi* only associates with a subset of *T. fuscocinerea*-like organisms that are typically parasitized by the more broadly distributed and closely related *C. striata* complex. Thus, we provide evidence that *C. bentleyi* may consistently target specific genotypes of a single host species, representing one of the narrowest host ranges for any mycoheterotroph reported thus far. Certainly, we could stand to learn more about the specific host genotypes that *C. bentleyi* prefers, as this would greatly aid in characterization of the niche requirements of this vulnerable orchid.

### Genetic variation

ISSR banding patterns based on two primers reveal that there is genetic variation within and among *C. bentleyi* localities. Perhaps the most surprising finding is that individuals within each sampling locality are not simply genetic clones of one another, as might be predicted for a predominantly selfing organism with extremely low population density and highly fragmented habitat. The observation of several color form variants, and of morphological variation within populations, is in concordance with our genetic findings based on ISSR variation. Most of the genetic diversity is instead found within populations of *C. bentleyi*, among plants sampled in some instances only a few meters apart. In fact, we found no evidence for identical clones among individuals within populations, considering all 75 polymorphic ISSR loci. This means one of two things: (1) that individuals are indeed able to outcross via pollinator-mediated pollen dispersal on a regional scale, despite never having been observed to do so; or (2) that multiple, genetically distinct, asexual lineages are present within each population. Given the low likelihood of outcrossing in this species, which is nearly always observed to set seed before flowers are open even in the most open-flowered populations (Fig. 1), a number of possible alternative explanations exist for the observed pattern of genetic variation within populations.

First, and the most superficially obvious hypothesis, would be dispersal among sites via seeds. Orchids are well known to produce thousands of microscopic “dust seeds,” lacking endosperm, which likely evolved as a strategy to “shower” the local environment with propagules in order to increase the chances that one or a few of these come into direct contact with an appropriate fungal partner for germination and development, given the initially mycoheterotrophic lifestyle shared by all orchids (Leake, 1994; Rasmussen, 1995; Arditti and Ghani, 2000; Rasmussen et al., 2015). It is not inconceivable that some propagules may travel via wind or some other dispersing agent to other suitable habitats several kilometers away, or perhaps even farther. However, evidence from studies of terrestrial orchid seed dispersal indicates that most seeds land within a few meters at most from the source plant, and beyond ∼10 m there is effectively zero recruitment (e.g. Jaquemyn et al., 2007; Jersáková and Malinová, 2007; Brzosko et al., 2017).

A second alternative involves a paleoclimatic scale: the possibility of persistent polymorphism following glacial retreat and range shifts from a once more continuous, larger, ancestral distribution. It is plausible, given the extent of glacial ice sheets and well-studied range shifts in suitable forest habitat as a result of Pleistocene glacial cycles (i.e. spanning the last 10^3^-10^7^ years; Williams et al., 2004), that *C. bentleyi* and its sister species in southern Mexico, *C. involuta*, shared a common ancestral range, having evolved vicariously from an ancestral species occupying habitats somewhere in the unglaciated portion of southern North America, as has been demonstrated for many other taxa sharing similar contemporary range disjunctions (e.g. Manchester, 1999; Williams et al., 2004; Copenhaver-Perry et al., 2017). Vicariance, perhaps as recently as over the last few thousand years, may have left unsorted ancestral polymorphism in remnant populations (e.g. Jaramillo-Correa et al., 2009). Thus, what is observed within and among localities for *C. bentleyi* may be remnants of a once broader geographic range, larger population sizes, and higher levels of genetic diversity, and the evolution of selfing may be a more recent development. In fact, *C. bentleyi* and *C. involuta* have likely been separated over a relatively short time period, possibly having diverged as recently as the last few thousands of years (Barrett et al., 2018).

A third alternative hypothesis involves an anthropogenic scale: very recent population assemblage and admixture following recolonization of second growth forest habitats from localized regional refugia, post-logging, over the last 100 years. The distribution of *C. bentleyi* was unknown before the late 1990s. Likewise, the ancestral habitat preferences of *C. bentleyi* and its fungal hosts are currently unknown. However, it may be safe to assume that prior to the extensive logging, mining, and other drastic anthropogenic disturbances of the last few centuries, it had a less fragmented distribution, likely harboring higher total genetic diversity, as in many other forest dwelling species in the eastern USA (e.g. Baucom et al., 2005). The plant populations surviving today likely represent a fraction of the previous population size and associated genetic diversity. The possibility exists that current populations represent assemblages of remnant genotypes that once inhabited more genetically structured populations, and thus have become intermixed following colonization of newly formed, secondary forest habitats. If colonization of these new habitats occurred rapidly from multiple source “microrefugia”, in which plants had ancestrally become differentiated and possibly locally adapted, the possibility exists what we observe today may be secondary contact/admixture zones.

A fourth possibility could be a lack of strong population-level resolution or high levels of homoplasy in putatively neutral ISSR markers, as is sometimes observed in SSR markers such as microsatellites (Garza and Freimer, 1996; Estoup et al., 2002). Despite our conservative and repeatable protocol for scoring bands (i.e. using bioanalyzer traces to confirm bands in agarose and polyacrylamide gels), the possibility exists that banding patterns exhibit homoplasy and obscure an underlying signal of population structure, e.g., as might be observed with SNP data. Further, the dominant nature of ISSR presence/absence banding reduces power to detect intra-individual variation among allelic variants for a particular locus. We are currently developing improvements to protocols that involve sequencing thousands of loci across the genome using multiple ISSR primer amplicons (e.g. Suyama and Matsuki, 2015), in order to detect neutral and adaptive polymorphisms as codominant markers in *C. bentleyi*.

A fifth, and final possibility involves symbiotic specificity. The compatible fungal genotypes present in a particular locality may exert selection on recruitment, germination, and survival of specific *C. bentleyi* genotypes, as has been suggested by Taylor et al. (2004) in *Corallorhiza maculata* (Raf.) Raf. Further, the use of predominantly neutral markers may obscure the detection of adaptive polymorphisms that potentially underlie host specificity. Unfortunately we were not able to sample sufficient numbers of *C. bentleyi* individuals to quantitatively compare patterns of ISSR variation and fungal host genotypes, which would be necessary to detect patterns of strong genotype-genotype interactions among *C. bentleyi* and its fungal hosts, as has been done preliminarily in other species of *Corallorhiza* (*C. maculata, C. odontorhiza* (Willd.) Poir.*, C. wisteriana* Conrad*, C. striata;* Taylor et al., 2004; Freudenstein and Barrett, 2014; Barrett et al., 2010).

What are the benefits of genetic diversity in a predominantly selfing species? Based on ISSR polymorphisms, most of the variation in *C. bentleyi* is found within sampling localities. However, it is unclear whether this would necessarily be beneficial to the species, if indeed all populations are exclusively autogamous. Autogamy prevents gene flow via pollen, and thus prevents genetic recombination among individuals, denying the opportunity for new allelic combinations in the face of changing conditions (Jullien et al., 2019). Autogamy is observed among many species of *Corallorhiza* (e.g. *C. odontorhiza, C. trifida* Châtel., *C. maculata*, and even other members of the *C. striata* complex). Potential explanations would be the absence of a suitable pollinator, or that the benefits of outcrossing are minimal among individuals that are genetically similar (in a relative sense). Orchids are “broadcast seed dispersers” due to the need for seeds to come into contact with a specific fungal host in order to germinate and develop. Thus, the benefits of selfing may be a way of ensuring sufficient local seed dispersal and survival that outweigh any potential benefits of outcrossing.

Selfing can evolve rapidly, in a few generations as a response to drastic environmental changes or pollinator decline/absence (Cheptou, 2019). A species or population with a low frequency of selfing may be rapidly swept to fixation in the absence of a pollinator (Schoen et al., 1996; Moeller and Geber, 2005; Barrett, 2014; Nieuwenhuis and Immler, 2016). Perhaps genetically distinct, selfing lineages found in *C. bentleyi* represent multiple selfing genotypes derived from more diverse ancestral populations of outcrossers or with mixed reproductive modes. It has been hypothesized that the evolution of selfing may represent an evolutionary “dead end” (e.g. Stebbins, 1950; Freyman and Höhna, 2019), but conclusions on this hypothesis are still equivocal (e.g. Wright et al., 2013). On one hand, selfing may lead to the accumulation of deleterious mutations over time, i.e. Müller’s Ratchet, without sexual recombination to allow the swapping of alleles among individuals. On the other hand, selfing may lead to accelerated differentiation among lineages or populations via reproductive isolation, and thus increased rates of speciation (e.g. Wendt et al., 2002; Briscoe Runquist et al., 2016; Sramkó et al., 2019). Through evolutionary history, occurrences of outcrossers evolving self-compatibility are far more common than the reversions from selfing to outcrossing (Stebbins et al., 1974; Barrett et al., 1996). However, if selfing is controlled single genes or regulatory elements (a “master regulator” such as a transcription factor), it is not inconceivable that a selfing lineage could revert back to outcrossing, especially if reproductive mode is under epigenetic control (e.g. Ellison et al., 2015).

More interestingly but less well tested, is the hypothesis that selfing is a way of fixing multilocus genotypes that allow the orchids to target specific fungal host genotypes. The rationale for this hypothesis is that recombination may “break up” coadapted allelic complexes that potentially allow fine-tuning of fungal host specificity, and thus selective pressure to repress recombination via the evolution of selfing may be advantageous (Taylor et al., 2004; 2013). In future studies, it will be necessary to take a genomic approach, sampling thousands of SNPs across the genome to detect patterns of fixed variants and linkage disequilibrium that are associated with fungal host genotypes, and to compare fitness metrics among selfing vs. outcrossing members of the same species in relation to fungal host utilization.

## CONCLUSIONS

The IUCN ‘vulnerable’ status assigned to *C. bentleyi* is based entirely on census assessment from a single survey taken almost two decades ago (400 individuals counted in 2001). This does not consider genetic, mating-system, or symbiotic attributes of a species, which are of particular concern for orchid conservation (Fay, 2018; Gale et al., 2018). In contrast to other extremely restricted eastern North American plant species (Lackey, 2004; Fernando et al., 2015), repeated, census-based assessment of population size and stability of *C. bentleyi* has not been conducted. The lack of long term, or even repeated, surveys have left an incomplete picture of extant localities, and the dynamics of population size and demography across flowering seasons. The predominant, and likely obligate reproductive mode of selfing in this species decreases the effective population size drastically relative to the census population size, meaning that census-based approaches may be misleading. The limited number of compatible fungal hosts with which this species associates implies that ecological niche space may be severely restricted by that of a few *Tomentella* host genotypes and their geographic distributions, about which almost nothing is known. While the finding of genetic variation within populations was surprising, and may seem beneficial, the predominance of selfing may restrict the adaptive potential of *C. bentleyi* with respect to rapid changes in the environment due to anthropogenic disturbance, changing climatic conditions, and competition with invasive species. In particular, hog peanut (*Amphicarpaea bracteata* (L.) Fernald) and Japanese stiltgrass (*Microstegium vimineum* (Trin.) A. Camus) have invaded several sites where *C. bentleyi* was known to occur (C. Barrett, personal observation). The former is a native species, but becomes highly invasive due to disturbance, while the latter was introduced from Asia and is highly invasive. In recent visits to those sites over the last three years—in Bath and Giles Cos., VA—no individuals of *C. bentleyi* were observed. Although many individual mycoheterotrophs are known to persist belowground for up to several years without flowering (Rasmussen, 1995; Bentley, 2000), the possibility exists that disturbance and invasive species have altered soil ecosystem function such that *C. bentleyi* can no longer survive at these sites. Clearly, additional sustained monitoring and studies of soil dynamics at invaded sites are both needed to assess the effects of invasive species on the survival of *C. bentleyi* and other rare forest plants.

Based on low census population sizes, restricted range, selfing (and its consequences for effective population size), extreme fungal host specificity (restriction of niche space), and encroachment of native habitats by invasive species, it is clear that the conservation status of *C. bentleyi* deserves further study. We advocate for regular, agency-led monitoring of *C. bentleyi* populations, including: multi-year census among focal localities to quantify fluctuations in the number of individuals flowering, and assessment of habitat quality (e.g. soil nutrients, tree diversity, abundance, stand age, co-occurring understory species richness and abundance, suitable fungal host availability, and assessment of threats from invasive species, e.g. Japanese stiltgrass). Ex-situ conservation could be conducted using seed packet methods (i.e. burying seeds in alternative locations in packets penetrable by fungal hyphae, *sensu* Rasmussen and Whigham, 1993) to encourage germination and growth in new habitats with seed collected from known localities. Alternatively, the transfer of tree seedlings or saplings in association with suitable fungal hosts could be conducted, followed by seed inoculations *ex situ*. Restrictions on anthropogenic disturbance in the vicinity of known occurrences of *C. bentleyi* (e.g. logging, mining operations, development) could improve or at least prevent further habitat degradation. Sustained mitigation and control of invasive species such as Japanese stiltgrass at known localities of *C. bentleyi*, which have the potential to negatively impact the number of flowering shoots and subsequent seed dispersal, though more study is needed on the specific impacts of invasives on the specific niche requirements of this orchid.

## Supporting information

Online Resources

## ACKNOWLEDGEMENTS

We thank Stanley Bentley, Charles Garratt, Nancy Van Alstine, John Freudenstein, and Zachary Bradford for field, technical, and photographic assistance; Ryan Percifield and Ashley Henderson for technical assistance with sequencing at the WVU Genomics Core Facility; and the USDA Forest Service for permission to collect samples. Funding was provided by NSF-EPSCOR (Award 1920858), the WVU Department of Biology, the WVU-Program to Stimulate Competitive Research, and the West Virginia Research Challenge Fund through a grant from the Division of Science and Research, HEPC and in part by the WVU Provost’s Office and WVU Eberly College of Arts and Sciences.

## ONLINE RESOURCES

**Online Resource 1** Trimmed alignment of all 1452 fungal ITS sequences from genus *Tomentella*

**Online Resource 2 A** Sample polyacrylamide gel image demonstrating repeatability of banding patterns for ISSR locus 868 among four individuals of C. bentleyi. **B** Bioanalyzer traces for three individuals (primer 868) overlaid. Y-axis = standardized band intensity; X-axis = standardized band length distribution. Arrow = band is present in two individuals (423B and 423C), and absent in the third individual (423A)

**Online Resource 3** Presence/absence matrix of ISSR bands, scored via Bioanalyzer traces and polyacrylamide or agarose gels (primers 868 + 873 concatenated). Loci are ordered by size, and those loci present in >95% or less than 5% of individuals were removed

## Notes

### Competing Interest Statement

The authors have declared no competing interest.

