## Supplementary material for "Integrating genetics, morphology, and fungal host specificity in conservation studies of a vulnerable, selfing, mycoheterotrophic orchid": Online Resources: Fig_S2_poly_gel_ISSR.pdf

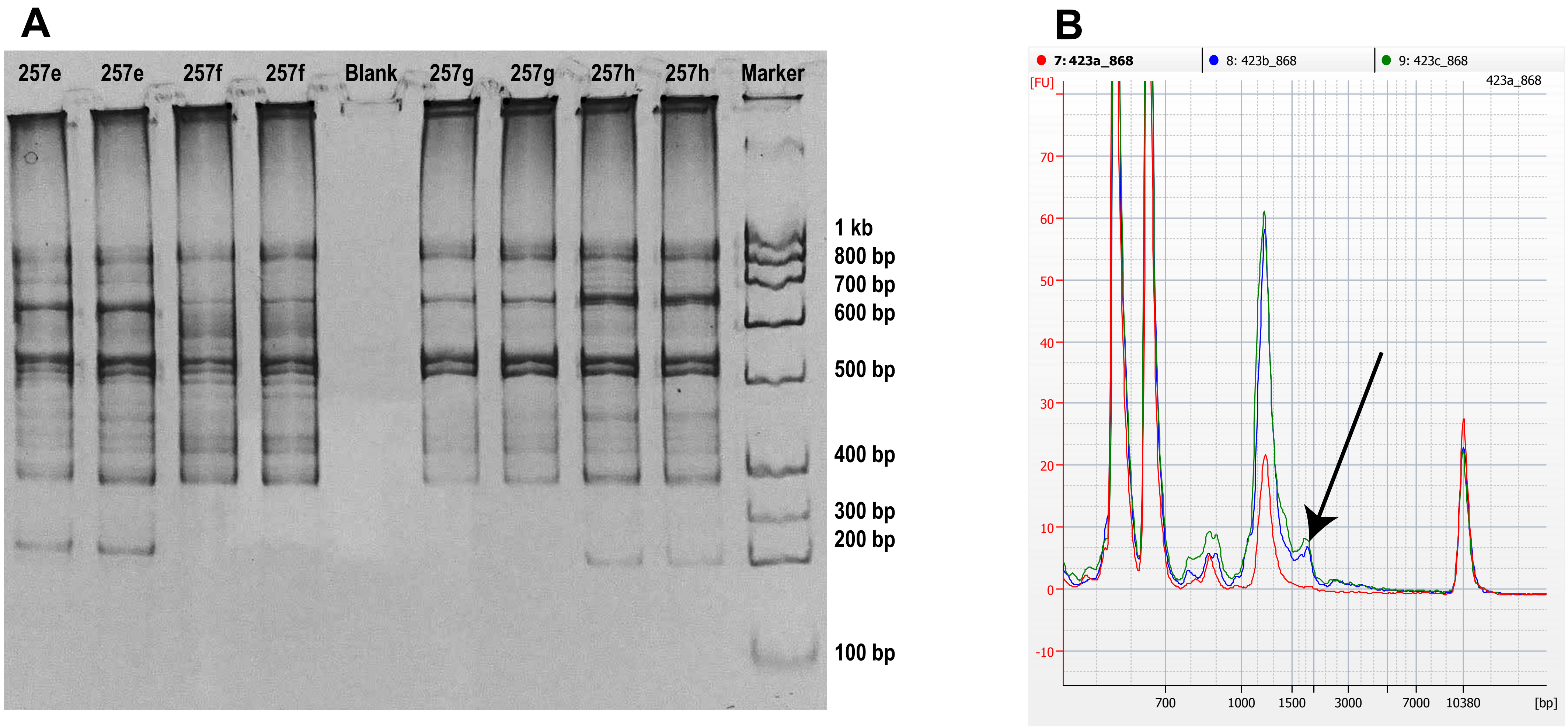


**Integrating genetics, morphology, and fungal host specificity in conservation assessment of a vulnerable, selfing, mycoheterotrophic orchid.** *Conservation Genetics*. Nicole M. Fama, Brandon T. Sinn, and Craig F. Barrett*.
Department of Biology, West Virginia University, 53 Campus Drive, Morgantown, West Virginia, USA 26506.
